# CircEML1 facilitates the steroid synthesis in follicular granulosa cells of chicken through sponging gga-miR-449a to release IGF2BP3 expression

**DOI:** 10.1101/2022.04.25.489339

**Authors:** Jing Li, Sujin Si, Xing Wu, Zihao Zhang, Chong Li, Yiqing Tao, Pengkun Yang, Donghua Li, Guoxi Li, Xiaojun Liu, Yadong Tian, Xiangtao Kang

## Abstract

Non-coding RNAs (ncRNAs) induced competing endogenous RNAs (ceRNA) play crucial roles in various biological process by regulating target gene expression. However, the studies of ceRNA networks in the regulation of ovarian ovulation process of chicken remains deficient compared to that in mammals. In the present study, it was revealed that circEML1 was differential expressed in hen’s ovarian tissue at different ages (15W, 20W, 30W and 68W) and identified as a loop structure from EML1 pre-mRNA, which promoted the expressions of CYP19A1 and StAR and the production of E2 and P4 in follicular granulosa cells (GCs) using qRT-PCR and ELISA. Furthermore, circEML1 was proved to serve as a sponge of gga-miR-449a to participate in the steroidogenesis using the dual luciferase reporter, RNA FISH assays, qRT-PCR and ELISA assays. In addition, we evaluated several potential target genes of gga-miR-449a and found that IGF2BP3 was targeted by gga-miR-449a and promoted steroidogenesis and E2/P4 secretions in GCs, which may act the regulatory role via mTOR/p38MAPK pathways. Meanwhile, we implemented a rescue experiment and demonstrated that gga-miR-449a reversed the promoting role of circEML1 on IGF2BP3 expression and steroidogenesis. Eventually, this study suggested that circEML1/gga-miR-449a/IGF2BP3 axis exerted an important role in the regulation of steroidogenesis and steroid hormones’ production possibly through mTOR/p38MAPK pathways in follicular GCs of chicken and may contribute a better understanding of ceRNA network in the modulatory mechanism of hen’s ovarian development and ovulation cycle.

## Introduction

In the past decade, a plenty of functional non-coding RNAs (ncRNAs), including microRNAs (miRNAs), long non-coding RNAs (lncRNAs) and circular RNAs (circRNAs) were identified within eukaryotic cells to mediate cellular biological processing through pre- and post-transcriptional regulation and then participate in the multiple regulating mechanisms of growth development, reproductive processing, even disease and cancer, and so on [1–4].In particular, miRNAs are well-known with a short sequence in 20-22 nt to suppress the transcriptional and translational levels of target genes typically by complementary pairing between the converted seed sequence of miRNA and the 3’-untranslated region (3’-UTR) of messenger RNA (mRNA)[5–7]. Recently, there were numerous studies have reported that the functional miRNAs play a primary role in the regulation of ovarian function, including follicular development and atresia, follicular granulosa cells (GCs) proliferation and apoptosis, steroid synthesis, and ovarian cancer[8–13], such as in mammals, miR-26b, miR-34a, miR-125a-5p, miR-182 and miR-92a could induce follicular granulosa cells (GCs) apoptosis by targeting their own genes’ 3’UTR [14–17], and miR-125b, miR-375, miR-150, miR-873 and miR-202 could influence steroid synthesis via inhibiting the expression of target genes[12, 18–20]; while in chicken, there were only several researches demonstrated that miR-26a-5p, miR-1b-2p, miR-205b and miR-23b-3p may be involved in the modulation of follicular development and steroid hormones synthesis[21–23]. Furthermore, circRNAs are generated from back-splicing of pre-mRNA as loop structures without 5’ to 3’ polarity and polyadenylated tail, including exonic circRNA, intronic circRNA and exonintron circRNA, which are highly stable in multiple tissues, and exonic circRNA have been reported to serve as a sponge of miRNA to affect the transcription and translation of downstream genes [24–26], while other types of circRNAs also exerted their functions via directly binding and encoding protein[27, 28]. Until now, numerous studies have proved that circRNAs could regulate mammals’ ovarian granulosa cell proliferation and apoptosis, steroidogenesis, as well as participate in the development and progression of ovarian cancer through sponging miRNAs and promoting the expression of downstream genes as a competing endogenous RNA (ceRNA) network, such as circRNA FGFR3/miR-29a-3p/E2F1 axis, circRNA UBAP2/miR-382-5p/PRPF8 axis, circRNA Cdr1as/miR-1270/SCAI axis, circWHSC1/miR-145 and miR-1182/ MUC1 and hTERT axis, and so on [29–32]. However, circRNAs act as a sponge of miRNA to regulate follicular development and steroid synthesis in chicken is still relatively lagging behind.

In our pervious study, we found serval potential functional miRNAs, especially gga-miR-449 family may be involved in the modulation of steroid synthesis, in the transcriptome sequencing analysis of ovarian tissue at the four different ages (15w, 20w, 30w and 68w) of hens [33]. Within a deeper research, we found that gga-miR-449a (5’-U**GGCAGUG**UAU GUUAGCUGGU -3’), gga-miR-449b-5p (5’-A**GGCAGUG**UGCUGUUAGCGGCUG -3’) and gga-miR-449c-5p (5’ U**GGCAGUG**CGUCUUAGCUGGCUGU 3’) has the same binding site in the gga-miR-449 family and found novel_circ_0010840 (circEML1) was predicted to have highly binding capacities with the three miRNAs using RNAhybird (http://bibiserv.techfak.uni-bielefeld.de/rnahybrid/), which obtained from our previous whole transcriptome sequencing dataset. Therefore, in this present study, we would investigate and seek the regulatory role of their possible regulatory ceRNA networks on steroid synthesis and hormones secretion in hen’s GCs, to reveal the functions of the non-coding RNAs on hens’ ovulation cycle and reproductive mechanism and provide a new approach for hens’ breeding and the improvement of egg-laying traits.

## Results

### CircEML1 promote the steroid synthesis and secretion of GCs

From the previous transcriptome database, novel_circ_0010840 was predicted to share the complementary sequence with gga-miR-449a, gga-miR449b-5p, and gga-miR-449c-5p and differently expressed in ovarian tissue at the age of 15W, 20W, 30W and 68W, especially highest expression presented in the ovary of 30W using the analysis of transcriptome and qRT-PCR, which might participate to the regulation of steroidogenesis in follicular GCs (Fig. 1a). Then, we identified the loop structure and distribution of novel_circ_0010840 using the PCR amplification, gel electrophoresis, RNase R treatment and the nuclear mass separation analysis. In the results, it demonstrated that novel_circ_0010840 (named as circEML1) originated from back-spliced exon 2, 3 and 4 circularization of EMAP like 1 (EML1) gene and had a highly stable expression in ovarian tissue of hens by treating RNase R (Fig. 1a-c) and a higher abundance in cytoplasmic of GCs compared to that in the nuclear (Fig. 1d), which indicated that circEML1 may function as a sponge of miRNA. Then, we evaluated the regulation of circEML1 on steroid synthesis and secretion in follicular GCs of hens by the qRT-PCR and ELISA assay and found that the expression of circEML1 was significantly promoted by treating circEML1 overexpression, which could notably increase the mRNA expressions of CYP19A1 and StAR (*p* < 0.01 or *p* < 0.05), while remarkably downregulate CYP11A1 expression in GCs (*p* < 0.05) (Fig. 1e and f). Furthermore, the concentrations of E2 and P4 in GC had a measurable rise after the stimulation of overexpression of circEML1 (*p* < 0.05 or *p* < 0.01), except for androgen’s level (Fig. 1g). Thus, all of these results indicated that circEML1 could facilitate steroid synthesis and the production of E2 and P4.

**Fig. 1.**
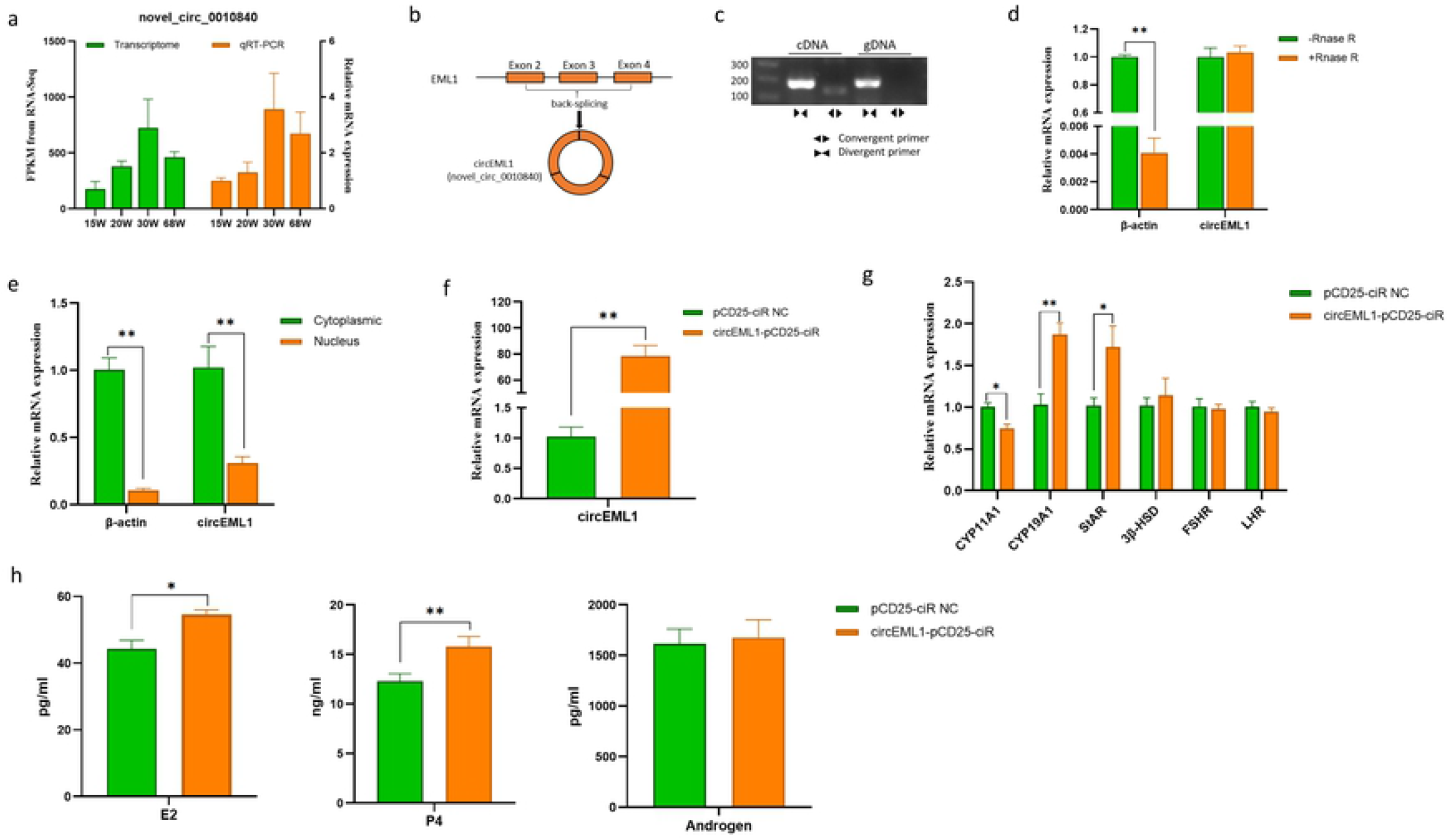
The expression and characteristics of circEML1 and the regulatory role of circEML1 on steroid synthesis in follicular GCs. (a) The relative expression of circEML1 in ovarian tissue at different ages of hens; (b)The loop structure of ciecEML1; (c) Divergent primers amplify circEML1 in cDNA but not genomic DNA (gDNA); (d) The expression of circEML1 and β-actin in ovarian tissue by treating RNase R; (e) The expression of circEML1 and β-actin in nucleus and cytoplasmic in granulosa cell; (f) The efficiency of circEML1 overexpression in GCs; (g) The relative mRNA expression of the key genes related steroid synthesis in GCs; (h) The concentrations of E2, P4 and androgen in GCs.

### CircEML1 acted as a sponge of gga-miR-449a

To deeply research the potential ceRNA networks of circEML1 with gga-miR-449a, gga-miR-449b-5p and gga-miR-449c-5p, we determined the expressions of the three miRNAs after transfected circEML1 overexpression in GCs and it was showed that circEML1 suppressed the expression of gga-miR-449a and gga-miR-449b-5p (*p* < 0.05), while it had no effect on the expression of gga-miR-449c-5p (Fig. 2a). Furthermore, we detected the target relationship with gga-miR-449a/gga-miR-449b-5p using the dual luciferase reporter assay, which displayed that the relative luciferase activity significantly decreased in the treatment of circEML1 WT and gga-miR-449a mimic compared to that in the other treatments, which indicated that circEML1 could directly bind to gga-miR-449a (*p* < 0.01) (Fig. 2b). However, there were no change in all the treatments when cotransfected with gga-miR-449b-5p mimic/NC and circEML1 WT/MUT. Moreover, FISH assay demonstrated that there was co-location of circEML1 and gga-miR-449a in the cytoplasmic (Fig. 2c). These results indicated that circRML1 could sponge gga-miR-449a and might facilitate the steroidogenesis of follicular GCs by downregulating its expression.

**Fig. 2.**
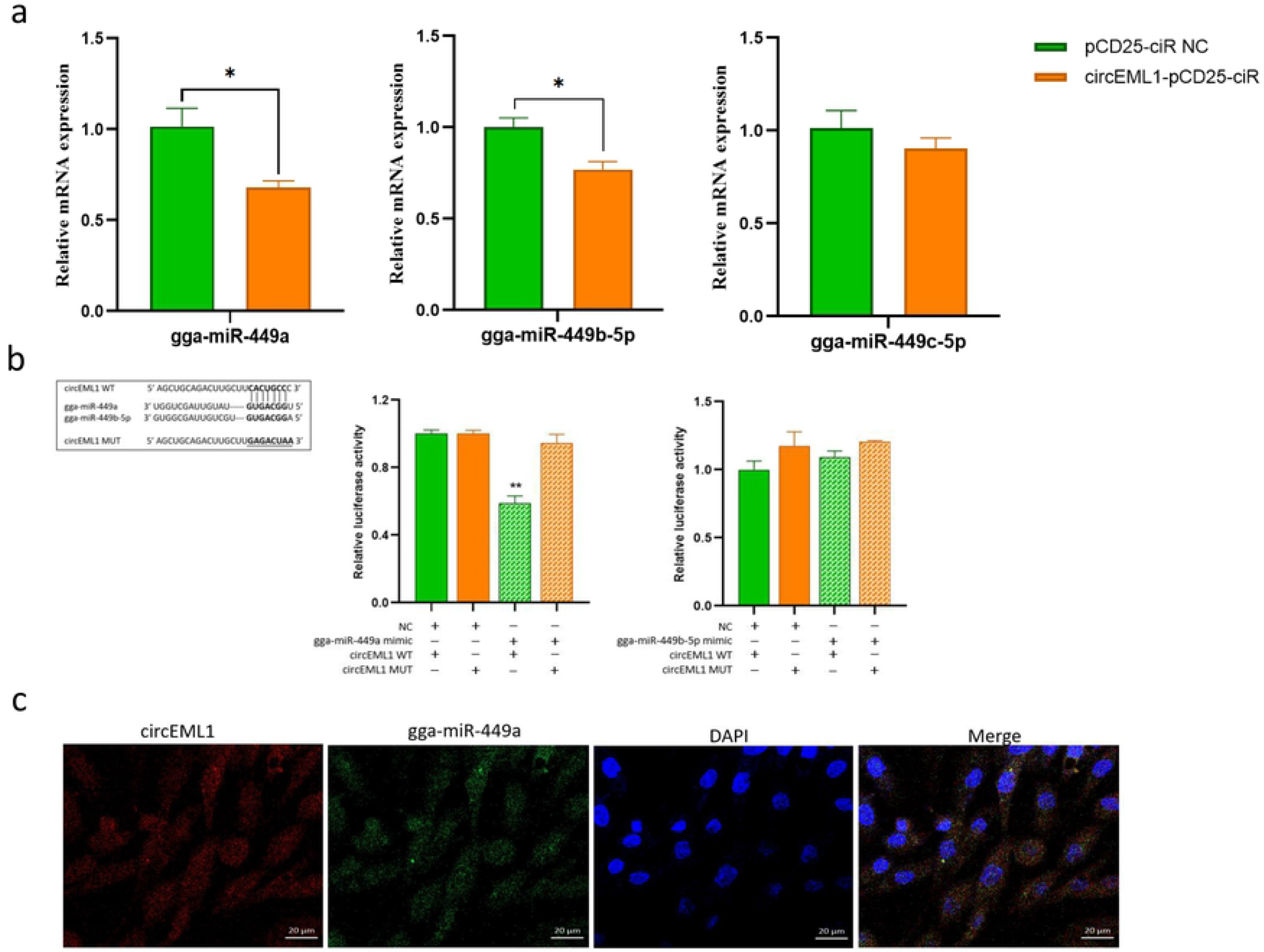
CircEML1 acted as a sponge of gga-miR-449a. (a) The relative expression of gga-miR-449a, gga-miR-449b-5p and gga-miR-449c-5p in GCs by treating circEML1 expression; (b) Dual luciferase reporter assay for detecting the interaction between circEML1 and gga-miR-449a/gga-miR-449b-5p in DF-1 cell; (c) RNA FISH assay for circEML1 and gga-miR-449a. Red, circEML1; Green, gga-miR-449a; Blue, DAPI; Merge, the merge of circEML1, gga-miR-449a and DAPI.

Then, to confirm the postulation, we transfected the mimic or inhibitor of gga-miR-449a into the GCs. The significant upregulation of gga-miR-449a was noticed after the stimulation of gga-miR-449a mimic, and vice versa (*p* < 0.01) (Fig. 3a). In addition, gga-miR-449a could alter the promotion of circEML1 on steroidogenesis and suppress the mRNA expression of CYP19A1 and StAR (*p* < 0.01), whereas upregulate the mRNA expression of CYP11A1 (*p* < 0.01), and inhibit the production of E2 and P4 in follicular GCs (*p* < 0.05) (Fig. 3b and c). These results indicated that circEML1 could modulate the steroid synthesis and secretion by sponging gga-miR-449a.

**Fig. 3.**
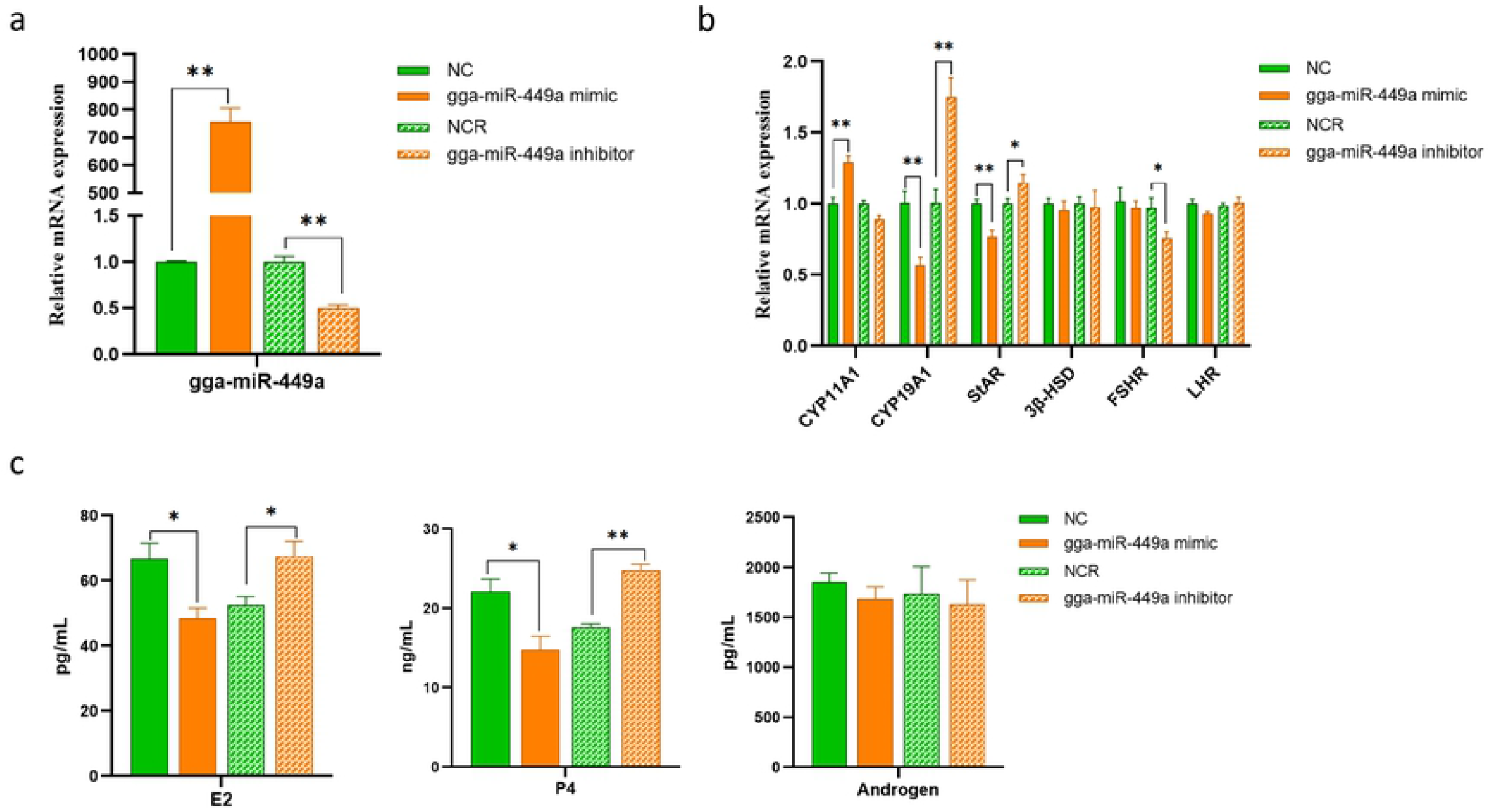
The effect of gga-miR-449a on the gene expression of the key genes related steroid synthesis and the secretion on steroid hormones. (a) The transfection efficiency of gga-miR-449a mimic or inhibitor; (b) The mRNA expression of key genes related steroid synthesis in GCs by transfecting gga-miR-449a; (c) The concentrations of steroid hormones in GCs by transfecting gga-miR-449a,

### IGF2BP3 was targeted by gga-miR-449a and promoted steroidogenesis in GCs

To better understand the action of circEML1 induced ceRNA network, we predicted several potential targeting gene associated with the modulatory role of gga-miR-449a on steroidogenesis by using TargetScan (http://www.targetscan.org/) and miRanda (http://www.microRNA.org/) database, including PGRMC1, MMP2, IGF2BP3, BMP3 and E2F5. Then, we detected the expression of these potential target genes after transfection with gga-miR-449a mimic. The results were demonstrated that gga-miR-449a expression was extremely higher in the treatment of gga-miR-449a mimic than that in control and the mRNA expression of IGF2BP3 could be significantly downregulated (*p* < 0.01), while the other potential genes were not impacted by gga-miR-449a (Fig. 4a and b). In addition, the protein expression of IGF2BP3 was also notably declined by treating gga-miR-449a mimic (*p* < 0.05) (Fig. 4c), all of which indicated that gga-miR-449a could target and inhibit the expression of IGF2BP3 at transcription and translation. Furthermore, the dual luciferase reporter system was conducted to further explore the target relationship of IGF2BP3 with gga-miR-449a and noticed that the luciferase activity of IGF2BP3 WT transfected with gga-miR-449a mimic in DF-1 cell were significantly decreased (*p*<0.01) compared to that in the other treatments, which indicated that IGF2BP3 3’UTR had the complementary binding site with gga-miR-449a (Fig. 4d). Then, we further identified the regulatory role of IGF2BP3 on steroid synthesis and hormones secretion in GCs and it demonstrated that overexpression of IGF2BP3 could extremely rise the mRNA and protein expression of IGF2BP3, and vice versa (Fig. 4e). Moreover, IGF2BP3 could upregulate the expression of CYP19A1 and StAR (*p* < 0.01 and *p* < 0.05), while downregulate the expression of CYP11A1 (*p* < 0.01), and increase the secretion of E2 and P4 (*p* < 0.01) (Fig. 4 f and g), which were consistent with the overexpression of circEML1 and opposite with the stimulation of gga-miR-449a. All of these results indicated that circEML1 facilitated the steroid synthesis and secretion in follicular GCs through the gga-miR-449a/IGF2BP3 axis.

**Fig. 4.**
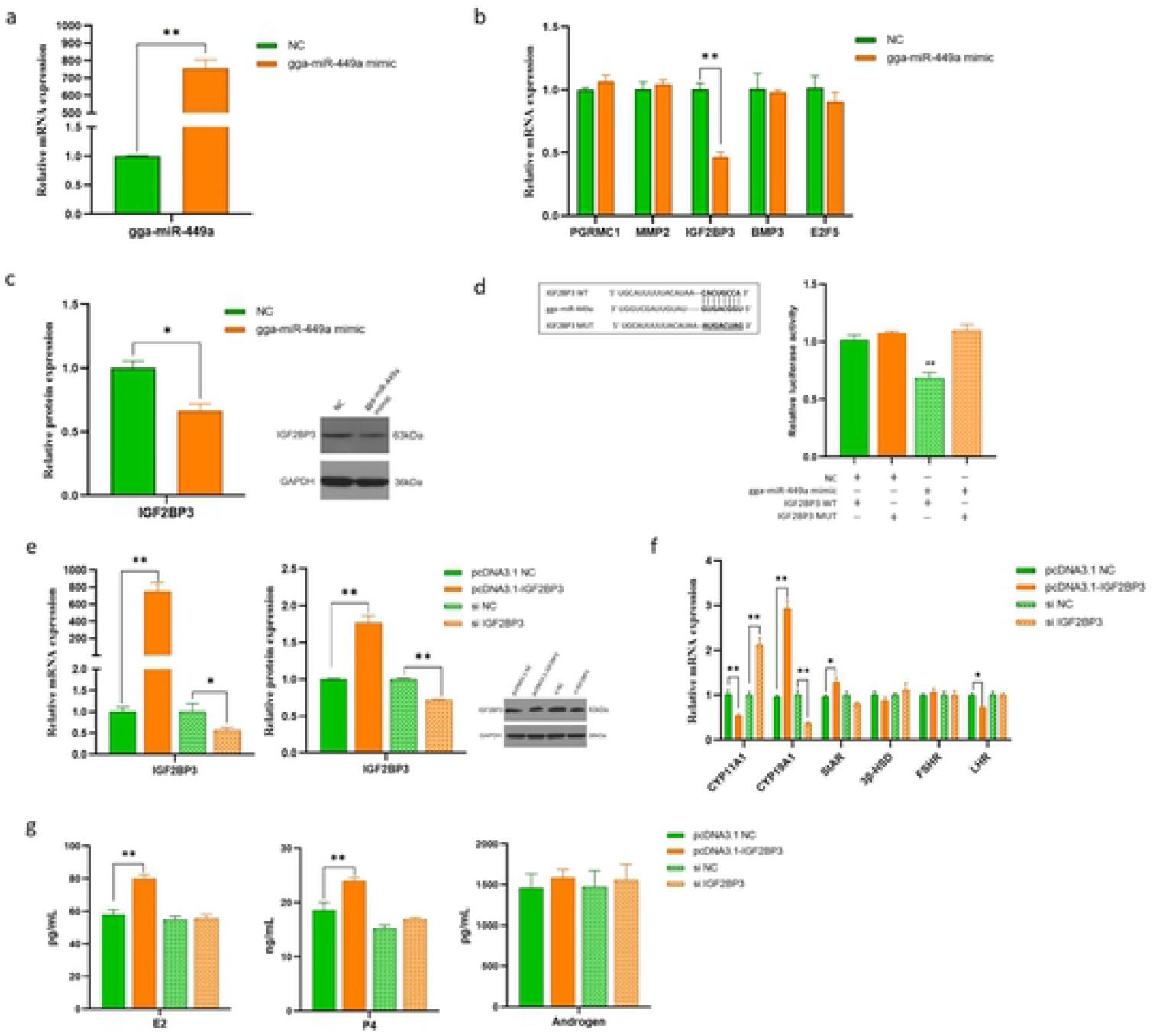
IGF2BP3 was targeted by gga-miR-449a and promoted the steroid synthesis and secretion. (a) The transfection efficiency of gga-miR-449a mimic; (b) The relative expression of potential target genes of gga-miR-449a in GCs after transfection with gga-miR-449a mimic; (c) The protein expression of IGF2BP3 in GCs after transfection with gga-miR-449a mimic; (d) The relative luciferase activity of the co-transfection of gga-miR-449a and IGF2BP3 in DF-1 cell; (e) The mRNA and protein expressions of IGF2BP3 in GCs after treating IGF2BP3 overexpression or interference; (f) The mRNA expression of key genes related steroid synthesis after transfecting IGF2BP3 overexpression or interference; (g) The concentrations of steroid hormones after transfecting IGF2BP3 overexpression or interference.

According to the above results, to search the further regulatory mechanism of IGF2BP3, we deeply studied the potential related signaling pathways of IGF2BP3 and treated with their inhibitors-AKT pathway (AKT-I), P13K pathway (PI3K-I), mTOR pathway (mTOR-I), p38MAPK pathway (p38MAPK-I) and IGF pathway (IGF-I), which have been reported before [34–36]. The results demonstrated that the expressions of IGF2BP3 have no significant difference among all the treatments expect to co-treat with overexpression of IGF2BP3, which revealed that IGF2BP3 expression wasn’t impacted by inhibiting these pathways (Fig. 5a). Moreover, the expressions of CYP19A1, StAR, CYP11A1 and 3β-HSD were significantly higher or lower (p < 0.01 or p < 0.05), and the levels of E2 and P4 have remarkably risen (p < 0.05) after co-transfected DMSO and overexpression IGF2BP3 compared to the treatment of DMSO alone in GC (Fig. 5a and b), all of which were generally consistent with the results of overexpression of IGF2BP3 alone. Some of the steroidogenesis genes’ expressions, the receptors’ expressions of FSH and LH, and the concentrations of E2 and P4 were significantly impacted after co-transfected the inhibitors of AKT, PI3K and IGF pathways with overexpression of IGF2BP3, separately, compared to those in the treatments of the above inhibitors alone (p < 0.01 or p < 0.05), which showed similar tendencies in comparison to those of the treatments of DMSO and DEMSO + IGF2BP3 (Fig. 5a and b). However, most of the key related genes of steroidogenesis and steroid hormones haven’t been changed between the treatments of mTOR-I and mTOR-I + IGF2BP3, or the treatments of p38MAPK-I and p38MAPK-I + IGF2BP3, which indicated that the regulatory role of IGF2P3 in steroid synthesis was suppressed if inhibiting mTOR or p38MAPK signaling pathways.

**Fig. 5.**
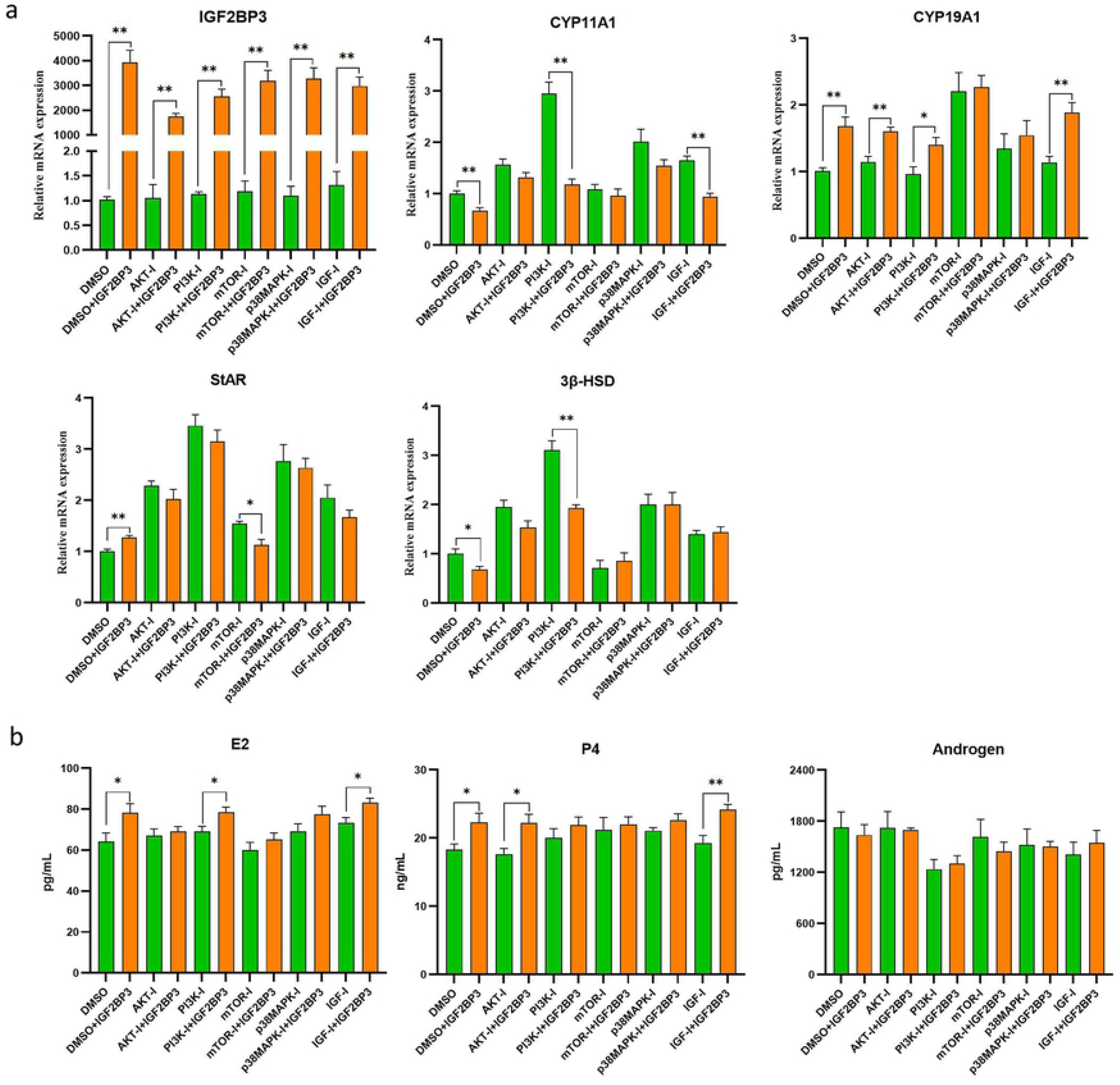
The evaluation of the potential related signaling pathways of IGF2BP3 regulating steroid hormone synthesis and hormone secretion. (a) The gene expression of IGF2BP3 and the key genes related steroid synthesis in GC after transfected the inhibitors of potential signaling pathway alone or co-transfected with overexpression of IGF2BP3; (b) The concentrations of steroid hormones in GC after transfected the inhibitors of potential signaling pathway alone or co-transfected with overexpression of IGF2BP3.

### Re-validation of the ceRNA network on the regulation of steroidogenesis

To re-valid circEML1 act a ceRNA to sponge gga-miR-449a and enhance the expression of IGF2BP3 and modulate the steroidogenesis in GCs, we performed a rescue experiment with cotransfecting circEML1 overexpression and gga-miR-449a mimic to investigate whether gga-miR-449a mimic can reverse the function of circEML1 or not. In the results, it was revealed that gga-miR-449a mimic could reverse the suppression of circEML1 in gga-miR-449a expression and the promotion in the mRNA and protein expression of downstream target gene-IGF2BP3 (*p* < 0.01 and *p* < 0.05) (Fig. 6a). Furthermore, it also eliminated the facilitation of circEML1 in the expression of CYP19A1 and StAR (*p* < 0.01 and *p* < 0.05), as well as the productions of E2 and P4 (*p* < 0.01) (Fig. 6b and c). Collectively, all of the above results suggested that circEML1 could upregulate the expressions of CYP19A1 and StAR and increase the secretions of E2/P4 in follicular GCs of hens by sponging gga-miR-449a to promote the mRNA and protein of IGF2BP3, which may play the regulatory role through activating mTOR/p38MAPK pathways (Fig. 7).

**Fig. 6.**
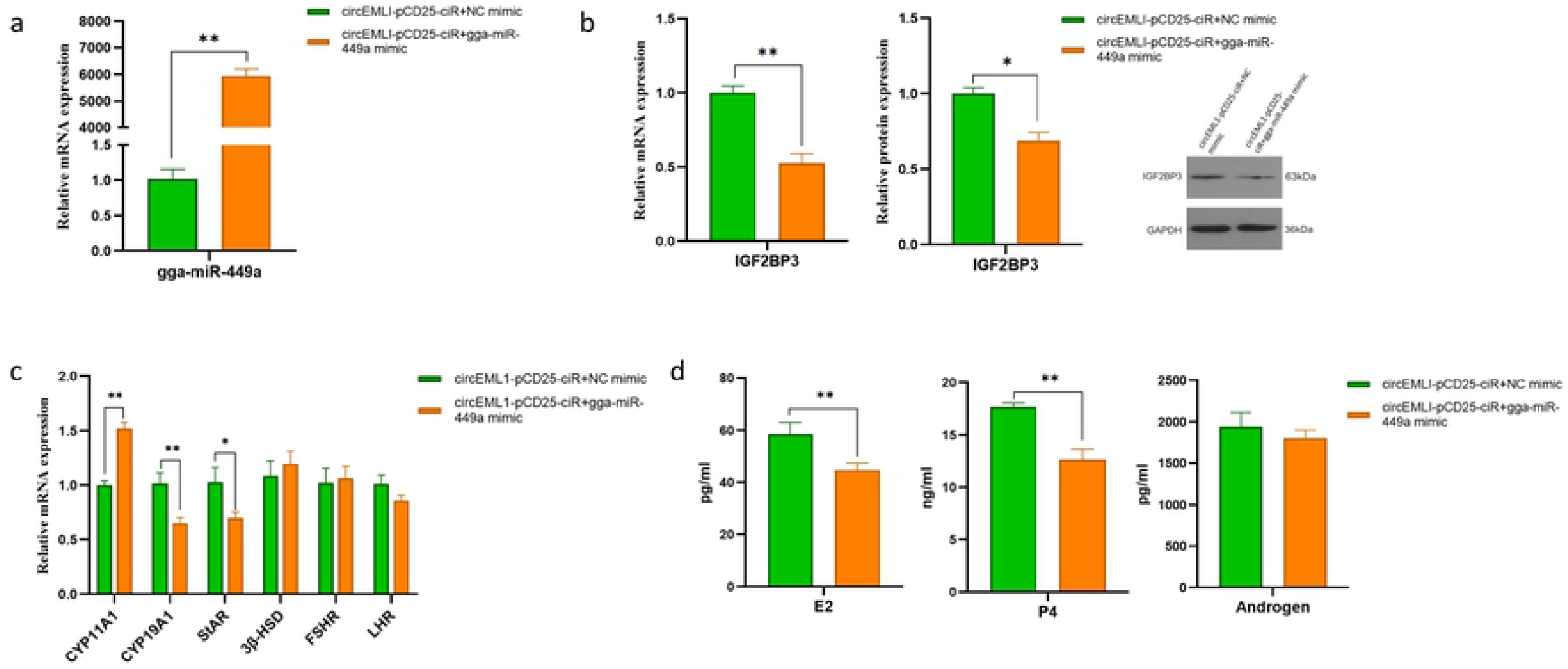
Gga-miR-449a reversed the regulatory role of circEML1 on the expressions of gga-miR-449a and IGF2BP3 and the promoting effect of steroid synthesis. (a) The mRNA expression of gga-miR-449a after transfection with cicrEML1 overexpression and gga-miR-449a mimic/NC in GCs; (b) The relative mRNA and protein expression of IGF2BP3 in GCs; (c) The relative mRNA expressions of the key genes related steroid synthesis in GCs; (d) The concentrations of E2, P4 and androgen in GCs.

**Fig. 7.**
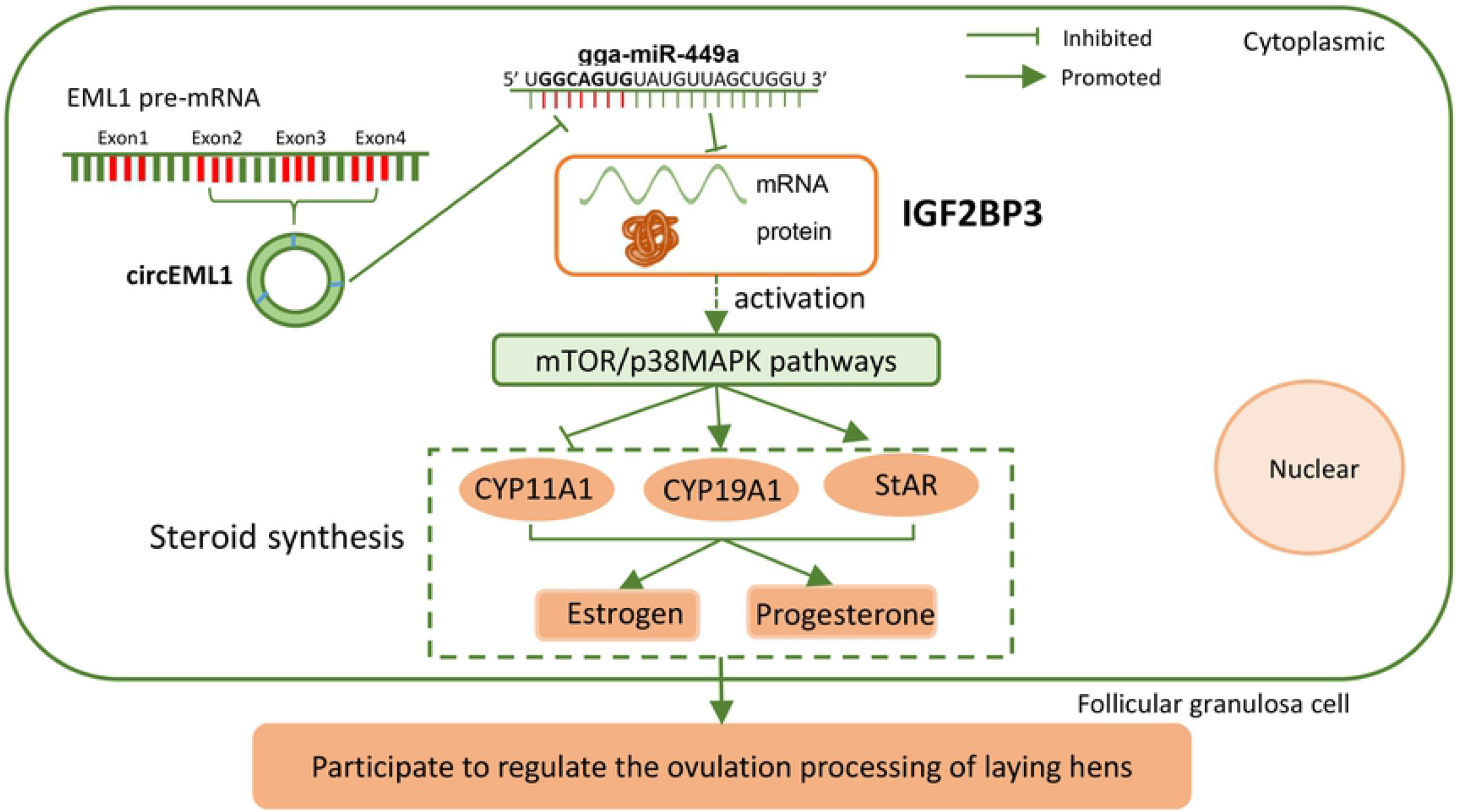
The working model of circEML1-induced ceRNA network.

## Discussion

Ovarian development and function were highly associated with the egg-laying performance of hens, and steroidogenesis and steroid hormones are essential for the whole ovulation processing of hens, including follicular selection, growth and maturation, even atrophy, and mainly estradiol and progesterone produced by follicular cells could stimulate follicular development and ovulation, leading to maintain egg-laying mechanism [37–39]. In our previous transcriptome database, we found circEML1 was differentially expressed in hens’ ovarian tissue at the ages of 15W, 20W, 30W and 68W, and the highest expression of circEML1 was presented in the ovary of 30W. It is well-known that hen’s ovary contained a large numerous of follicles with different sizes and strongly produced steroid hormones to maintain the ovulation processing during the peak laying period (30-40w), which indicated that the regulatory role of circEML1 might be associated with steroidogenesis[40, 41]. In recent years, circRNAs have been proven to drive as a sponge of miRNA to competitively bind miRNA and release the expression of downstream target genes, which participated in the modulation of reproductive mechanism in various mammals, including follicular GC proliferation, apoptosis and steroidogenesis, for instance, circEGFR increased E2 secretion and enhance ovarian GC proliferation by regulating miR-125a-3p and Fyn; ssc-circINHA-001 regulated follicular GC apoptosis via sponging miR-241-5p, miR-7144-3p and miR-9830-5p to enhance the expression of downstream gene INHBA; and the depletion of circLDLR could decrease estrogen secretion via regulating miR-1294 and CYP19A1 expression as a ceRNA network [42–44]. However, there were only few researches of circRNA reported to regulate the ovarian development in avian, such as circRNA aplacirc_013267 could promote granulosa cell apoptosis by sponging apla-miR-1-13 and increasing the expression of THBS1 [45]. Therefore, in the present study, we investigated the regulatory role of circEML1 on steroid synthesis and secretion, and discovered that circEML1 could facilitate the expressions of CYP19A1 and StAR, as well as the productions of E2 and P4, as we expected. In addition, we predicted a high binding ability of circEML1with gga-miR-449a, gga-miR-449b-5p and gga-miR-449c-5p (shared a same seed sequence) and further verified that circEML1 acted as a sponge of gga-miR-449a in cytoplasm to suppress the expression of gga-miR-449a.

Interestingly, in mammals, the two clusters of miR-449a/b/c family and miR-34b/c family were known to have overlapping functionality in multiple biological processes, including the male reproductive system and ovarian cancer’s development [46–49]. Furthermore, circGFRA1, circCCNDB1 have been discovered to inhibit cell senescence and promote ovarian cancer progression by sponging miR-449a, respectively [50]. However, in our study, gga-miR-449a was proven to inhibit the steroid synthesis and steroid hormones’ secretions in follicular granulosa cells of chicken, which was opposite with the effect of circEML1 and indicated that circEML1 could function as gga-miR-449a sponge to regulate the steroidogenesis of hen’s GCs. In addition, we further investigated several potential target genes to improve the regulatory mechanism of circEML1/gga-miR-449a-induced ceRNA network and found that gga-miR-449a could target IGF2BP3 and inhibited the mRNA and protein expression of IGF2BP3 in follicular GCs. However, IGF2BP3 is well-known as a member of insulin-like growth factor-2 mRNA-binding proteins, and promote target mRNAs stability and translation as a N6-methyladenosine (m^6^A) reader, such as IGF2, MYC, ACTB and AML, which is also present in multiple tissues and mainly associated with tumorigenesis and tumor development as a biomarker of cancer [51–56]. In recent two years, there were a few miRNAs have been reported to target IGF2BP3 for the regulation of different tissue and cancer cells proliferation and invasion, such as mi-127-5p, miR-9-5p, miR-370, miR-142-5p, miR-320b and so on [57–61]. Although there was no direct report of IGF2BP3 participated in ovarian steroid synthesis, some studies have reported IGF2BP3 modulated early embryo and oocyte development in aqua [62, 63]. Furthermore, numerous researches have demonstrated that insulinlike growth factor 2 (IGF2) can be involved in endocrine and paracrine regulation of mammals’ ovarian function, including follicular development, follicular atresia and the promotion of steroidogenesis and steroid hormones excretion, which was post-transcriptionally regulated by IGF2BP3[39, 64–66], and all of those implied that IGF2BP3 might exert a major role on the ovarian function of chicken. Thus, our results demonstrated that overexpression of IGF2BP3 could upregulate the expression of CYP19A1 and StAR and facilitate the levels of E2 and P4 in follicular GCs, which was consistent with those results of overexpression of circEML1 and inhibition of gga-miR-449a. Some researches have reported that IGF2BP3 could act as an upstream regulator of AKT/mTOR signaling in hepatocellular carcinoma and promote nasopharyngeal carcinoma migration and invasion by activating the AKT/mTOR signaling pathway[36, 67]. In addition, IGF2BP3 has also been revealed to evoke p53, p38/MAPK signal transduction events, and involved in the occurrence and development of acute myeloid leukemia (AML)[68]. However, in this study, we found that the regulatory function of IGF2BP3 on steroid synthesis and hormones secretion depressed if co-treated with the inhibitors of mTOR and p38MAPK pathways in follicular GC, which indicated that the regulation of IGF2BP3 may act through the activation of mTOR/ p38MAPK pathways and enriched the biological functions of IGF2BP3.

Furthermore, we performed a rescue experiment to re-valid the function of ceRNA network through overexpression of circEML1 and gga-miR-449a mimic/NC. It was demonstrated that gga-miR-449a could reverse the modulatory function of circEML1 on the mRNA and protein expression of IGF2BP3 and steroid synthesis and E2/P4 secretions. Therefore, all of these results have provided a robust evidence that cirEML1 could facilitate steroid synthesis in follicular GCs of chicken by sponging miR-449a to promote the expression of downstream target gene-IGF2BP3.

Collectively, this present study first systematically illustrated that circEML1-induced ceRNA network (circEML1/gga-miR-449a/IGF2BP3) could facilitate steroidogenesis and the secretions of E2 and P4 in follicular GCs of hens’ ovary by sponging gga-miR-449a to promote IGF2BP3 expression, which possibly play the regulatory role in follicular GCs by activating mTOR/p38MAPK signaling pathways. Meanwhile, these findings may contribute a better understanding of non-coding RNAs’ function on steroid synthesis and ovarian ovulation process, as well as provide a new approach for laying hens’ breeding and the improvement of egg-laying traits.

## Materials and Methods

### Ethics approval

The animals used in this study were treated in accordance with the guide for the administration of affairs concerning experimental animals and approved by the Animal Care and Use Committee at Henan Agricultural University (Permit Number: 19-0068).

### Granulosa cell collecting and culture

A total of 40 laying hens at the ages of 30~40 weeks-old were chosen to remove the whole ovary tissue, then stored in a PBS buffer with adding 3% double antibody and immediately isolated the granulosa layer from prehierarchical follicles (Lovell and T., 2003). All of the collected granulosa cells were mixed into an RNAase-free 1.5 ml centrifuge tube and rapidly cut into small pieces, then added a 0.25% trypsin with equal volume and digested for 10mins in an incubator at 37°C and 5% CO_2_. The obtained solution was filtered using a 200μm screen and the precipitates were collected by centrifugation at 1,800 rpm for 5mins. Then we received cell suspension using the complete medium with 2.5% bovine serum and 1% double antibody after repeating the centrifugation process when suspended the precipitates with a PBS solution. The cell suspension was plated in 12-well plates and cultured in an incubator with the same setting as above for growing ~12 h before transfection, as described in the previous study [33].

### Characterization and distribution of circEML1

The convergent and divergent primers of circEML1 were designed by Oligo 7 and synthesized by Qingke Bio (Shown in Table 1). The cDNA and gDNA samples were extracted and obtained from the above hens, and then the cDNA and gDNA PCR products were run on 2% agarose gel electrophoresis after PCR amplification treatment. RNase R assay was carried out by incubating total RNA of ovarian tissue with or without RNase R treatment at 37°C for 30min. The treated RNA was reverse transcribed into cDNA and determined the expression of circEML1 by qRT-PCR.

**Table 1.**
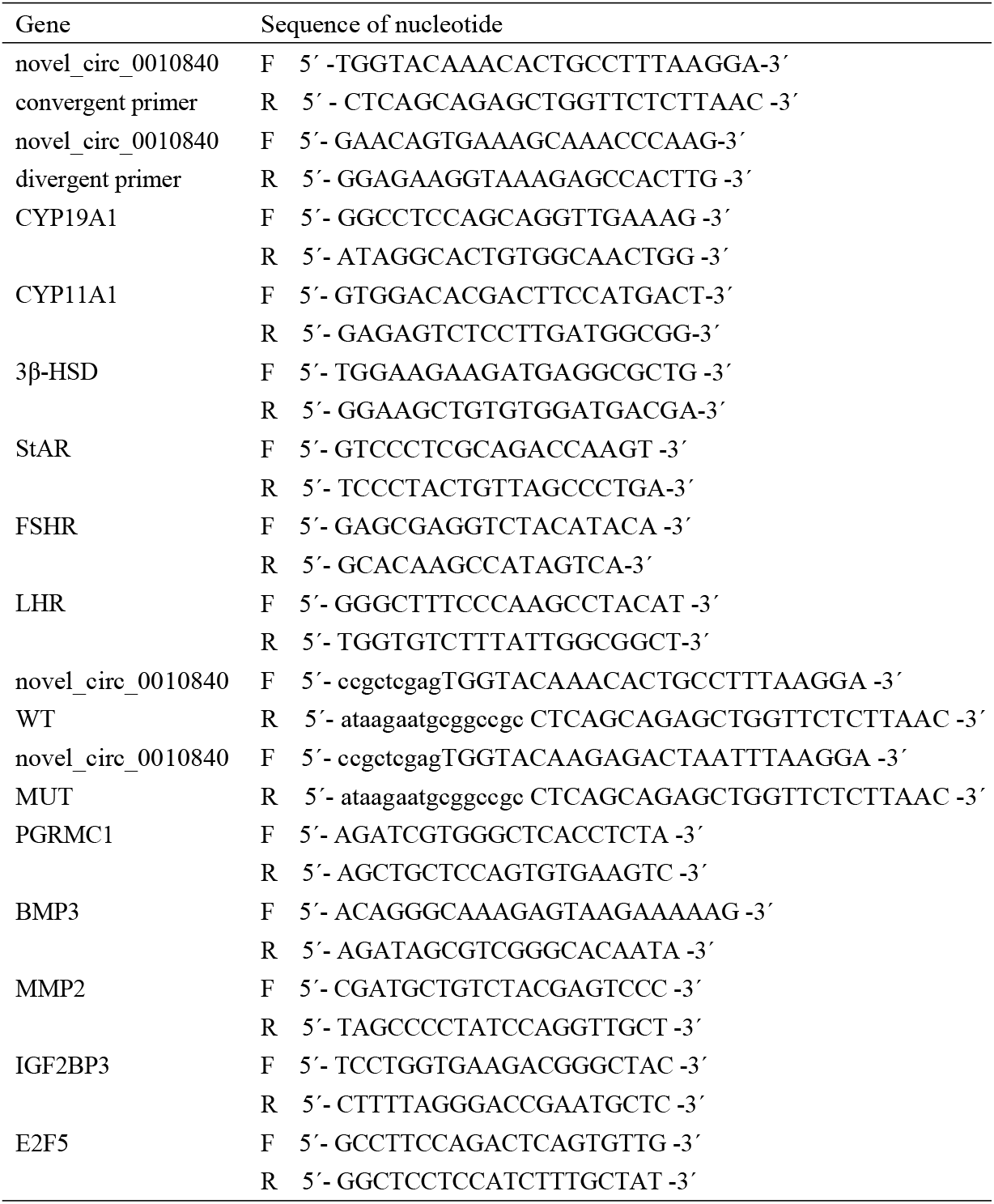
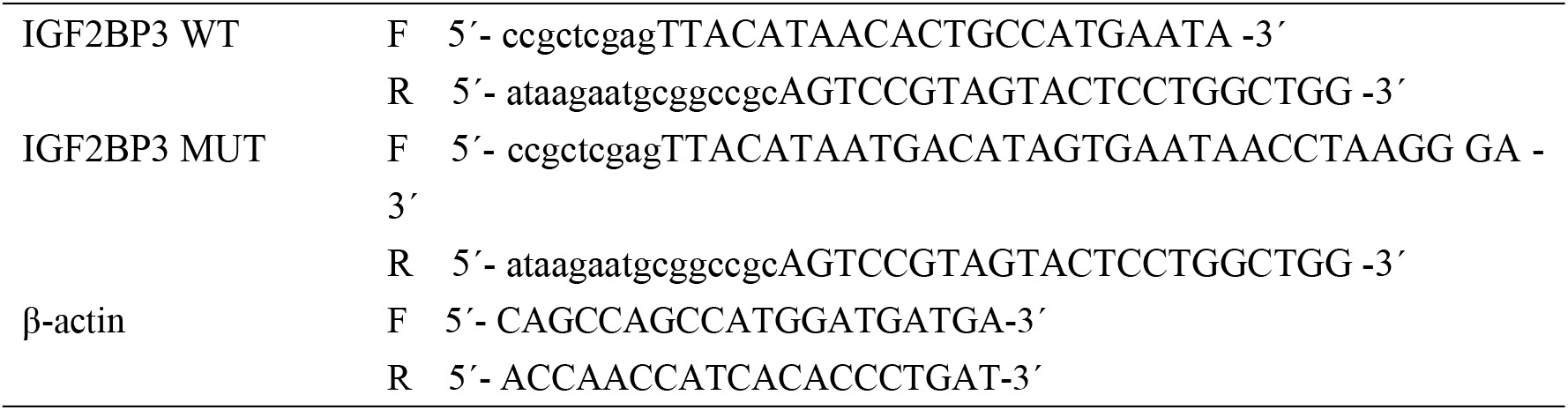
The primers of genes presented in the study

GCs were cultured and digested by 0.25% trypsin before transfection, then collected into an RNAase-free 1.5 ml centrifuge tube for the distribution of circEML1 in cell nucleus and cytoplasm assay. The isolation and extraction of GCs nucleus and cytoplasm fractions were performed using PAPIS^tm^ Kit (Invitrogen, Shanghai, China) following the manufacturer’s protocols and detected the expression of circEML1 by qRT-PCR.

### Plasmids construction and cell transfection

The overexpression plasmid of novel_circ_0010840 were synthesized by Geneseed Biotech (Guangzhou, China), the full-length was cloned into pCD25-ciR vector to obtain circEML1-pCD25-ciR, and pCD25-ciR was used as its negative control (pCD25-ciR NC). The mimics and inhibitors of gga-miR-449a and their negative controls (NC and NCR) were synthesized by RiboBio (Guangzhou, China). The overexpression plasmid of IGF2BP3 (pcDNA3.1-IGF2BP3) were built and cloned by isolating chicken IGF2BP3 CDS region and inserting into pcDNA3.1-EGFP vector using restriction enzymes *EcoRI* and *HindIII*, and the cloned pcDNA3.1-EGFP was used as a negative control (pcDNA 3.1 NC). The interference fragment-IGF2BP3 (si IGF2BP3) and its negative control (si NC) were purchased from RiboBio (Guangzhou, China). The follicular GCs were cultured into 12-well cell plates and transfected with the treatment of pCD-ciR NC, circEML1-pCD25-ciR, gga-miR-449a mimic/NC, gga-miR-449a inhibitor/NCR, pcDNA3.1 NC, pcDNA3.1-IGF2BP3, si NC, si IGF2BP3, circ EML1-pCD25-ciR +NC mimic or circEML1-pCD25-ciR+gga-miR-449a mimic after the cell density reached >70% using Lipofectamin™ 3000 or riboFECT CP Transfection Kit, individually, according to the manufacturer’s protocols. Furthermore, the inhibitors of AKT (1μm Afuresertib hydrochloride), PI3K(25μm LY294002), mTOR(100μm Rapamycin), p38MAPK(20μm SB203580), and IGF(20μm BMS536924) pathways were selected with 0.1% DMSO as negative control, which were transfected alone or co-transfected with overexpression of IGF2BP3 in follicular GC using LipofectaminTM 3000 Kit (Invitrogen, Shanghai China), respectively. After that, cells were incubated for 36h and collected for subsequent experiments.

### Quantitative real time PCR (qRT-PCR)

Total RNA of GC was extracted using Trizol reagent after 36h post-transfection, and then frozen at −80 °C for the following study. The gene and miRNA cDNA was obtained from RNA using HiScript^®^ III 1st Strand cDNA Synthesis Kit (+gDNA wiper) and HiScript® III RT SuperMix for qRCR (+gDNA wiper) (Vazyme, Nanjing, China). The quantitative real time PCR (qRT-PCR) was performed to determine the relative expressions of genes and miRNAs using ChamQ Universal SYBR qPCR Master Mix (Vazyme, Nanjing, China) according to the manufacturer’s guideline. And the relative expressions of genes and miRNAs were calculated using the 2^-ΔΔCt^ method with β-actin and U6 as their reference genes, respectively. All the primers including the following primers used in this study were designed by Oligo 7 software and listed in Table 1.

### ELISA assay for steroid hormones

The cell supernatant per well was extracted after centrifugation 3000 rpm for 20min and determined steroid hormones’ concentrations according to the guidelines of the chicken Estradiol (E2), progesterone 4 (P4) and Androgen ELISA kits, which provided by Jiangsu Meimian Industrial Co., Ltd (Yangcheng, China).

### West-blotting assay

The qRT-PCR assay was performed to determine the mRNA expression of IGF2BP3 after post transfection in follicular GC following the process in above. The west-blotting assay was carried out to determine the protein concentration of IGF2BP3 in follicular GC. The protein of GC were collected after 36h post-transfection and treated using Pierce TM Rapid Gold BCA protein assay kit (Elabscience, Wuhan, China) according to the manufacturer’s protocols. The primary antibodies of IGF2BP3 and GAPDH were purchased from Novusbio (Littleton, CO, USA) and Affinity (Cincinnati, OH, USA), respectively, and the secondary antibody (HRP-Goat anti Rabbit) was provided by Elabscience (Wuhan, China). The OD value of target band was detected by the AlphaEaseFC software system.

### Dual-luciferase reporter assay

The wild type (WT) and mutant type (MUT) primers of the sequence of circEML1 and the 3’UTR region of IGF2BP3 were designed to contain or change the predicted binding site with gga-miR-449a or gga-miR-449b-5p (shown in Table 1), and cloned into psiCHECK II plasmid with restriction enzymes *Not* I and Xhol I (Takara, Dalian, China). The obtained plasmids were named as circEML1 WT, circEML1 MUT, IGF2BP3 WT and IGF2BP3 MUT, individually.

DF-1 cell, as a well-known cell line of chicken embryo fibroblasts, was used to validate the gga-miR-449a/gga-miR-449b-5p target interactions with circEML1/IGF2BP3. The cells were incubated in a complete medium with DMEM (F12), 10% bovine serum and 1% double antibody, and then seeded into 12-well plates for incubation and co-transfected circEML1 WT, circEML1 MUT, IGF2BP3 WT or IGF2BP3 MUT with gga-miR-449a mimic, gga-miR-449b-5p mimic or NC using Lipofectamin™ 3000 (Invitrogen, Shanghai China) after the cell density reached >90%, respectively.

The cells per well were washed with a PBS solution and completely lysed with 100μl 1xPLB after 36h post-transfection, and then the supernatant was extracted by centrifugation at 7,500 rpm for 1 min and a 20μl loaded into a well of 96-well plate with adding 100μl luciferase assay reagent (LARΠ) and 100μl Stop&Glo reagent in turn according to the guideline of the Dual Luciferase Reporter Assay System Kit (Promega, Madison, WI, USA). The determination of luciferase activities was performed using Gen5™ microplate reader and imager software (BioTek, Vermont, USA).

### RNA fluorescence in situ hybridization (FISH) assay

The RNA-FISH assay was performed to detect the subcelluar co-location of circEML1 and gga-miR-449a in DF-1 cell using the specific probes. The probes of circEML1 (5’-TGGTCTTGCACTGAACAT-3’) and gga-miR-449a (5’-ACCAGCTAACATCTGCCA-3’) was synthesized by Gene seed Biotech and the treated cells were detected using the laser confocal microscope (Leica TCS SP2 AOBS, Germany).

### Statistical analysis

All the data were presented as means ± SEM and subjected to statistical analysis by one-way ANOVA of IBM SPSS Statistics v22.0. The statistical significance between groups was evaluated by Tukey HSD analysis, where p<0.05 (*) was considered significant differences and p<0.01 (**) was considered extremely significant differences. The data analysis was performed using GraphPad Prism 8 software.

## Author Contributions

J Li, SJ Si, W Xing, ZH Zhang, C Li and YQ Tao performed the experimental process. J Li and SJ Si wrote the manuscript. DH Li and PL Y conducted all the data analysis. YD Tian, XT Kang, GX Li and XJ Liu contributed to the experimental design. All authors approved the final manuscript.

## Acknowledgments

The study was supported by Program for Innovative Research Team (in Science and Technology) in University of Henan Province (21IRTSTHN022), Zhongyuan Science and Technology Innovation Leading Scientist Project (214200510003), and China Agriculture Research System (CARS-40-K04). Additionally, the authors would like to thank Henan Innovative Engineering Research Center of Poultry Germplasm Resource for providing study materials and site, and team supporting for sample collection and analysis.

## Declarations

### Consent for publication

Not applicable.

### Abbreviations

NcRNA: Non-coding RNA
ceRNA: Competing endogenous RNA
GCs: Granular cell
IGF2BP3: Insulin like growth factor 2 mRNA binding protein 3
EML1: EMAP like 1
CYP19A1: Cytochrome P450 family 19 subfamily A member 1
CYP11A1: Cytochrome P450 family 11 subfamily A member 1
StAR: Steroidogenic acute regulatory protein
3β-HSD: 3 beta- and steroid delta-isomerase 2
FSHR: Follicle stimulating hormone receptor
LHR: Luteinizing hormone receptor
E2: Estrogen
P4: Progesterone
miRNA: microRNA
lncRNA: long noncoding RNA
circRNA: circular RNA
E2F1: E2F transcription factor 1
PRPF8: Pre-mRNA processing factor 8
SCAI: Suppressor of cancer cell invasion
MUC1: Mucin 1 cell surface associated
hTERT: human telomerase reverse transcriptase
PGRMC1: Progesterone receptor membrane component 1
MMP2: Matrix metallopeptidase 2
BMP3: Bone morphogenetic protein 3
E2F5: E2F transcription factor 5
INHBA: Inhibin subunit beta A
THBS1: Thrombospondin 1
FSH: Follicle stimulating hormone
LH: Luteinizing hormone
IGF2: Insulin like growth factor 2
MYC: MYC proto-oncogene
ACTB: Actin beta
AML: Acute myeloid leukemia

## References

1. Anastasiadou E, Jacob LS, Slack FJ. Non-coding RNA networks in cancer. Nature reviews. Cancer. 2018; 18(1):5–18.

2. Palazzo AF, Lee ES. Non-coding RNA: what is functional and what is junk? Frontiers in genetics. 2015; 6:2.

3. Mattick JS, Makunin IV. Non-coding RNA. Human molecular genetics. 2006; 15 Spec No 1:R17–29.

4. Esteller M. Non-coding RNAs in human disease. Nature reviews. Genetics. 2011; 12(12):861–874.

5. Grimson A, Farh KK, Johnston WK, Garrett-Engele P, Lim LP, Bartel DP. MicroRNA targeting specificity in mammals: determinants beyond seed pairing. Molecular cell. 2007; 27(1):91–105.

6. Buchan JR, Parker R. Molecular biology. The two faces of miRNA. Science (New York, N.Y.). 2007; 318(5858):1877–1878.

7. Kehl T, Backes C, Kern F, Fehlmann T, Ludwig N, Meese E et al. About miRNAs, miRNA seeds, target genes and target pathways. Oncotarget. 2017; 8(63):107167–107175.

8. Gebremedhn S, Salilew-Wondim D, Ahmad I, Sahadevan S, Hossain MM, Hoelker M et al. MicroRNA Expression Profile in Bovine Granulosa Cells of Preovulatory Dominant and Subordinate Follicles during the Late Follicular Phase of the Estrous Cycle. PloS one. 2015; 10(5):e0125912.

9. Yerushalmi GM, Salmon-Divon M, Ophir L, Yung Y, Baum M, Coticchio G et al. Characterization of the miRNA regulators of the human ovulatory cascade. Scientific reports. 2018; 8(1):15605.

10. Zhang J, Xu Y, Liu H, Pan Z. MicroRNAs in ovarian follicular atresia and granulosa cell apoptosis. Reproductive biology and endocrinology: RB&E. 2019; 17(1):9.

11. Worku T, Rehman ZU, Talpur HS, Bhattarai D, Ullah F, Malobi N et al. MicroRNAs: New Insight in Modulating Follicular Atresia: A Review. International journal of molecular sciences. 2017; 18(2).

12. Sen A, Prizant H, Light A, Biswas A, Hayes E, Lee HJ et al. Androgens regulate ovarian follicular development by increasing follicle stimulating hormone receptor and microRNA-125b expression. Proceedings of the National Academy of Sciences of the United States of America. 2014; 111(8):3008–3013.

13. Maalouf SW, Liu WS, Pate JL. MicroRNA in ovarian function. Cell and tissue research. 2016; 363(1):7–18.

14. Tu F, Pan ZX, Yao Y, Liu HL, Liu SR, Xie Z et al. miR-34a targets the inhibin beta B gene, promoting granulosa cell apoptosis in the porcine ovary. Genetics and molecular research: GMR. 2014; 13(2):2504–2512.

15. Wang C, Li D, Zhang S, Xing Y, Gao Y, Wu J. MicroRNA-125a-5p induces mouse granulosa cell apoptosis by targeting signal transducer and activator of transcription 3. Menopause (New York, N.Y.). 2016; 23(1):100–107.

16. Liu J, Du X, Zhou J, Pan Z, Liu H, Li Q. MicroRNA-26b functions as a proapoptotic factor in porcine follicular Granulosa cells by targeting Sma-and Mad-related protein 4. Biology of reproduction. 2014; 91(6):146.

17. Liu J, Yao W, Yao Y, Du X, Zhou J, Ma B et al. MiR-92a inhibits porcine ovarian granulosa cell apoptosis by targeting Smad7 gene. FEBS letters. 2014; 588(23):4497–4503.

18. Donadeu FX, Sontakke SD, Ioannidis J. MicroRNA indicators of follicular steroidogenesis. Reproduction, fertility, and development. 2016.

19. Yu C, Li M, Wang Y, Liu Y, Yan C, Pan J et al. MiR-375 Mediates CRH Signaling Pathway in Inhibiting E2 Synthesis in Porcine Ovary. Reproduction (Cambridge, England). 2016.

20. Geng XJ, Zhao DM, Mao GH, Tan L. MicroRNA-150 regulates steroidogenesis of mouse testicular Leydig cells by targeting STAR. Reproduction (Cambridge, England). 2017; 154(3):229–236.

21. Kang L, Yang C, Wu H, Chen Q, Huang L, Li X et al. miR-26a-5p Regulates TNRC6A Expression and Facilitates Theca Cell Proliferation in Chicken Ovarian Follicles. DNA and cell biology. 2017; 36(11):922–929.

22. Li J, Hou L, Sun Y, Xing J, Jiang Y, Kang L. Single nucleotide polymorphism rs737028527 (G>A) affect miR-1b-3p biogenesis and effects on chicken egg-laying traits. Animal reproduction science. 2020; 218:106476.

23. Wei Q, Li J, He H, Cao Y, Li D, Amevor FK et al. miR-23b-3p inhibits chicken granulosa cell proliferation and steroid hormone synthesis via targeting GDF9. Theriogenology. 2022; 177:84–93.

24. Hansen TB, Jensen TI, Clausen BH, Bramsen JB, Finsen B, Damgaard CK et al. Natural RNA circles function as efficient microRNA sponges. Nature. 2013; 495(7441):384–388.

25. Patop IL, Wüst S, Kadener S. Past, present, and future of circ RNA s. The EMBO journal. 2019; 38(16):e100836.

26. Chen L-L, Yang L. Regulation of circRNA biogenesis. RNA biology. 2015; 12(4):381–388.

27. Du WW, Fang L, Yang W, Wu N, Awan FM, Yang Z et al. Induction of tumor apoptosis through a circular RNA enhancing Foxo3 activity. Cell death and differentiation. 2017; 24(2):357–370.

28. Zheng X, Chen L, Zhou Y, Wang Q, Zheng Z, Xu B et al. A novel protein encoded by a circular RNA circPPP1R12A promotes tumor pathogenesis and metastasis of colon cancer via Hippo-YAP signaling. Molecular cancer. 2019; 18(1):47.

29. Zhou J, Dong ZN, Qiu BQ, Hu M, Liang XQ, Dai X et al. CircRNA FGFR3 induces epithelial-mesenchymal transition of ovarian cancer by regulating miR-29a-3p/E2F1 axis. Aging. 2020; 12(14):14080–14091.

30. Zhao Z, Ji M, Wang Q, He N, Li Y. Circular RNA Cdr1as Upregulates SCAI to Suppress Cisplatin Resistance in Ovarian Cancer via miR-1270 Suppression. Molecular therapy. Nucleic acids. 2019; 18:24–33.

31. Zong ZH, Du YP, Guan X, Chen S, Zhao Y. CircWHSC1 promotes ovarian cancer progression by regulating MUC1 and hTERT through sponging miR-145 and miR-1182. Journal of experimental & clinical cancer research: CR. 2019; 38(1):437.

32. Xu Q, Deng B, Li M, Chen Y, Zhuan L. circRNA-UBAP2 promotes the proliferation and inhibits apoptosis of ovarian cancer though miR-382-5p/PRPF8 axis. Journal of ovarian research. 2020; 13(1):81.

33. Li J, Ma XJ, Wu X, Si SJ, Li C, Yang PK et al. Adiponectin modulates steroid hormone secretion, granulosa cell proliferation and apoptosis via binding its receptors during hens’ high laying period. Poultry science. 2021; 100(7):101197.

34. Suvasini R, Shruti B, Thota B, Shinde SV, Friedmann-Morvinski D, Nawaz Z et al. Insulin growth factor-2 binding protein 3 (IGF2BP3) is a glioblastoma-specific marker that activates phosphatidylinositol 3-kinase/mitogen-activated protein kinase (PI3K/MAPK) pathways by modulating IGF-2. Journal of Biological Chemistry. 2011; 286(29):25882–25890.

35. Mancarella C, Scotlandi K. IGF2BP3 from physiology to cancer: novel discoveries, unsolved issues, and future perspectives. Frontiers in Cell and Developmental Biology. 2020:363.

36. Xu Y, Guo Z, Peng H, Guo L, Wang P. IGF2BP3 promotes cell metastasis and is associated with poor patient survival in nasopharyngeal carcinoma. Journal of cellular and molecular medicine. 2022; 26(2):410–421.

37. Proszkowiec-Weglarz M, Rzasa J, Słomczyńska M, Paczoska-Eliasiewicz H. Steroidogenic activity of chicken ovary during pause in egg laying. Reproductive biology. 2005; 5(2):205–225.

38. Navara KJ. The role of steroid hormones in the adjustment of primary sex ratio in birds: compiling the pieces of the puzzle. Integrative and comparative biology. 2013; 53(6):923–937.

39. Nielsen J, Christiansen J, Lykke-Andersen J, Johnsen AH, Wewer UM, Nielsen FC. A family of insulin-like growth factor II mRNA-binding proteins represses translation in late development. Molecular and cellular biology. 1999; 19(2):1262–1270.

40. Li J, Li C, Li Q, Li G, Li W, Li H et al. Novel regulatory factors in the hypothalamic-pituitary-ovarian axis of hens at four developmental stages. Frontiers in genetics. 2020:1367.

41. Kang W, Yun J, Seo D, Hong KC, Ko Y. Relationship among egg productivity, steroid hormones (progesterone and estradiol) and ovary in Korean Native Ogol Chicken. Asian-Australasian Journal of Animal Sciences. 2001; 14(7):922–928.

42. Huang X, Wu B, Chen M, Hong L, Kong P, Wei Z et al. Depletion of exosomal circLDLR in follicle fluid derepresses miR-1294 function and inhibits estradiol production via CYP19A1 in polycystic ovary syndrome. Aging. 2020; 12(15):15414–15435.

43. Huang X, Wu B, Chen M, Hong L, Kong P, Wei Z et al. Depletion of exosomal circLDLR in follicle fluid derepresses miR-1294 function and inhibits estradiol production via CYP19A1 in polycystic ovary syndrome. Aging (Albany NY). 2020; 12(15):15414.

44. Ma M, Wang H, Zhang Y, Zhang J, Liu J, Pan Z. circRNA-Mediated Inhibin–Activin Balance Regulation in Ovarian Granulosa Cell Apoptosis and Follicular Atresia. International journal of molecular sciences. 2021; 22(17):9113.

45. Wu Y, Xiao H, Pi J, Zhang H, Pan A, Pu Y et al. The circular RNA aplacirc_13267 upregulates duck granulosa cell apoptosis by the apla-miR-1-13/THBS1 signaling pathway. Journal of cellular physiology. 2020; 235(7-8):5750–5763.

46. Comazzetto S, Di Giacomo M, Rasmussen KD, Much C, Azzi C, Perlas E et al. Oligoasthenoteratozoospermia and infertility in mice deficient for miR-34b/c and miR-449 loci. PLoS genetics. 2014; 10(10):e1004597.

47. Loukas I, Skamnelou M, Tsaridou S, Bournaka S, Grigoriadis S, Taraviras S et al. Fine-tuning multiciliated cell differentiation at the post-transcriptional level: contribution of miR-34/449 family members. Biological reviews of the Cambridge Philosophical Society. 2021; 96(5):2321–2332.

48. Yuan S, Liu Y, Peng H, Tang C, Hennig GW, Wang Z et al. Motile cilia of the male reproductive system require miR-34/miR-449 for development and function to generate luminal turbulence. Proceedings of the National Academy of Sciences of the United States of America. 2019; 116(9):3584–3593.

49. Wu J, Bao J, Kim M, Yuan S, Tang C, Zheng H et al. Two miRNA clusters, miR-34b/c and miR-449, are essential for normal brain development, motile ciliogenesis, and spermatogenesis. Proceedings of the National Academy of Sciences of the United States of America. 2014; 111(28):E2851–2857.

50. Yu AQ, Wang ZX, Wu W, Chen KY, Yan SR, Mao ZB. Circular RNA CircCCNB1 sponges micro RNA-449a to inhibit cellular senescence by targeting CCNE2. Aging. 2019; 11(22):10220–10241.

51. Hammer NA, Hansen T, Byskov AG, Rajpert-De Meyts E, Grøndahl ML, Bredkjaer HE et al. Expression of IGF-II mRNA-binding proteins (IMPs) in gonads and testicular cancer. Reproduction (Cambridge, England). 2005; 130(2):203–212.

52. Bell JL, Wächter K, Mühleck B, Pazaitis N, Köhn M, Lederer M et al. Insulin-like growth factor 2 mRNA-binding proteins (IGF2BPs): post-transcriptional drivers of cancer progression? Cellular and molecular life sciences: CMLS. 2013; 70(15):2657–2675.

53. Mancarella C, Scotlandi K. IGF2BP3 From Physiology to Cancer: Novel Discoveries, Unsolved Issues, and Future Perspectives. Front Cell Dev Biol. 2019; 7:363.

54. Huang H, Weng H, Sun W, Qin X, Shi H, Wu H et al. Recognition of RNA N 6-methyladenosine by IGF2BP proteins enhances mRNA stability and translation. Nat Cell Biol. 2018; 20(3):285–295.

55. Zhang N, Deng J, Zhou F. IGF2BP3 As an N6-Methyladenosine Reader Promotes Acute Myeloid Leukemia Progression By Regulating the Stability of Target RNA. Blood. 2021; 138:4322.

56. Ren Y, Du D, Liu T, Wang C, Yan Q, Li C et al. The m6A reader IGF2BP3 Facilitates Gastric Cancer Progression Through Activating EMT via Upregulating MYC mRNA Stability. 2021.

57. Zhang W, Liu H, Jiang J, Yang Y, Wang W, Jia Z. CircRNA circFOXK2 facilitates oncogenesis in breast cancer via IGF2BP3/miR-370 axis. Aging. 2021; 13(14):18978–18992.

58. Zhang W, Zhu L, Yang G, Zhou B, Wang J, Qu X et al. Hsa_circ_0026134 expression promoted TRIM25-and IGF2BP3-mediated hepatocellular carcinoma cell proliferation and invasion via sponging miR-127-5p. Bioscience reports. 2020; 40(7).

59. Yin H, He H, Shen X, Zhao J, Cao X, Han S et al. miR-9-5p Inhibits Skeletal Muscle Satellite Cell Proliferation and Differentiation by Targeting IGF2BP3 through the IGF2-PI3K/Akt Signaling Pathway. International journal of molecular sciences. 2020; 21(5).

60. Liu J, Jiang X, Zou A, Mai Z, Huang Z, Sun L et al. circIGHG-Induced Epithelial-to-Mesenchymal Transition Promotes Oral Squamous Cell Carcinoma Progression via miR-142-5p/IGF2BP3 Signaling. Cancer research. 2021; 81(2):344–355.

61. Wu K, Wang X, Yu H, Yu Z, Wang D, Xu X. LINC00460 facilitated tongue squamous cell carcinoma progression via the miR-320b/IGF2BP3 axis. Oral diseases. 2021.

62. Ren F, Lin Q, Gong G, Du X, Dan H, Qin W et al. Igf2bp3 maintains maternal RNA stability and ensures early embryo development in zebrafish. Commun Biol. 2020; 3(1):94.

63. Wu X, Zhang Y, Xu S, Chang Y, Ye Y, Guo A et al. Loss of Gsdf leads to a dysregulation of Igf2bp3-mediated oocyte development in medaka. Gen Comp Endocrinol. 2019; 277:122–129.

64. Tkachenko OY, Wolf S, Lawson MS, Ting AY, Rodrigues JK, Xu F et al. Insulin-like growth factor 2 is produced by antral follicles and promotes preantral follicle development in macaques†. Biology of reproduction. 2021; 104(3):602–610.

65. Giudice LC. Insulin-like growth factor family in Graafian follicle development and function. Journal of the Society for Gynecologic Investigation. 2001; 8(1 Suppl Proceedings):S26–29.

66. Spicer LJ, Aad PY. Insulin-like growth factor (IGF) 2 stimulates steroidogenesis and mitosis of bovine granulosa cells through the IGF1 receptor: role of follicle-stimulating hormone and IGF2 receptor. Biology of reproduction. 2007; 77(1):18–27.

67. Chen C-L, Tsukamoto H, Liu J-C, Kashiwabara C, Feldman D, Sher L et al. Reciprocal regulation by TLR4 and TGF-β in tumor-initiating stem-like cells. The Journal of clinical investigation. 2013; 123(7):2832–2849.

68. Qin T, Cheng Y, Wang X. RNA-binding proteins as drivers of AML and novel therapeutic targets. Leukemia & Lymphoma. 2022:1–13.

